# Learning the wiring rules of a mammalian cortical column

**DOI:** 10.64898/2026.07.09.737432

**Authors:** Oren Richter, Elad Schneidman

## Abstract

Characterization of neural circuits’ architecture typically relies on measurable neuronal features such as morphology, molecular identity, and spatial location. While generative models leveraging these properties have proven accurate, they remain constrained by available measurements and our assumptions regarding the prospective features. Here, we present an alternative approach using representational learning and use it to model the circuitry of a column of the mouse primary visual cortex. Our framework learns jointly low-dimensional embeddings of neurons in an abstract feature space alongside wiring rules that predict synaptic connectivity. These embedding-based models accurately predict individual synapses, connectivity degrees, and network motif statistics — outperforming standard generative models that depend on detailed cell-type classifications — using only a handful of embedding dimensions and wiring rules. Crucially, the learned representations prove interpretable, recapitulating cortical depth, cell type, and dendritic morphology. The resulting wiring blueprint is both simple and biologically meaningful, suggesting that cortical connectivity follows surprisingly parsimonious logic. This framework offers a general and exportable tool for learning minimal generative models of connectomes.

## Introduction

Mapping the relations between the structure of neural circuits and their function is fundamental to our understanding of how information is represented and processed by the brain, and how neural circuits direct decisions and actions [1–7]. The richness of functions that neural circuits perform stems from the biophysical properties of individual neurons, the diverse developmental plans that underlie neuronal wiring [8– 11], and the learning algorithms that optimize the connectivity map between neurons [12–16]. Thus, identifying the design principles of biological neural networks is crucial for delineating their capacity to learn, their computational abilities, and how we might fix or augment them.

The reconstructions of complete connectivity maps of neural circuits at the resolution of single cells and synapses [17–26] makes the quantitative study of the design and function of biological neural networks possible [27]. Generative models offer a natural framework for identifying and characterizing the biological and physical features that govern the architecture of neural networks, especially considering their stochastic, developmental, and variable nature [28, 29]. Indeed, such models proved to be highly accurate in recapitulating observed connectivity patterns, using a surprisingly simple handcrafted set of biological and physical features and rules [30–32]. However, these models are inherently biased and potentially limited by the assumptions of their designers and by the neuronal features that have been experimentally measured. Critically, the structure of neural circuits may rely on neuronal features that we cannot immediately name, identify, or even measure.

We study here the architecture of the connectome of a cortical column [33, 34] using generative models that we train to predict the connection probabilities between all neurons in the circuit. Importantly, to avoid the biases and assumptions of neural connectivity models that use a predefined set of biological and physical features (Fig. 1A, left), we utilize a representational learning approach [35–38] and rely on connectivity data alone: We learn models of the cortical connectome by finding both an abstract set of features that describe each of the neurons, i.e. an “embedding” in an abstract space, and “wiring rules”, i.e. an Artificial Neural Network (ANN), that use these features to determine which neurons would be connected (Fig. 1A, right).

**Fig. 1.**
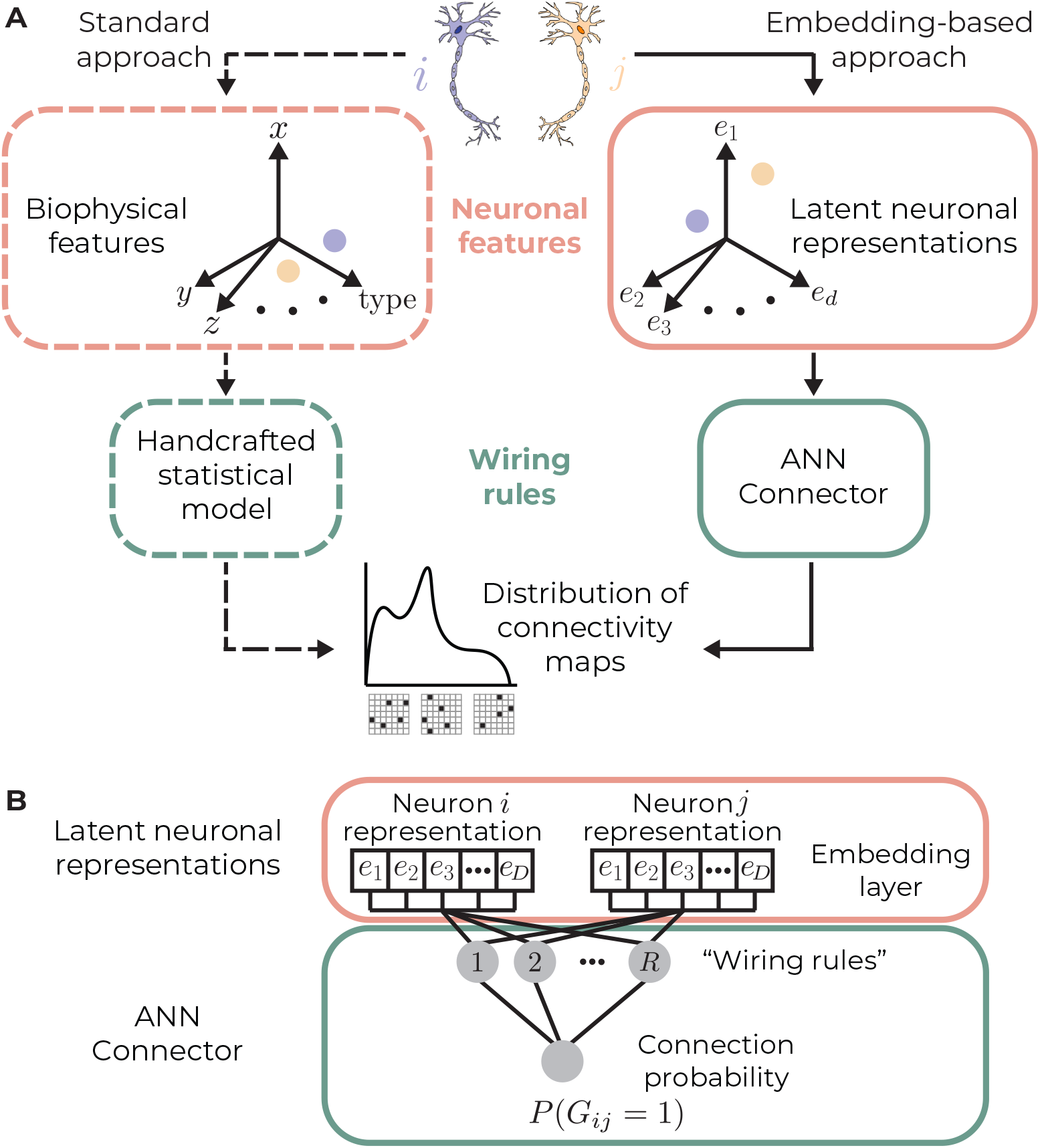
An embedding-based approach for learning neuronal features and wiring rules simultaneously. **A**. A cartoon comparing the “standard” approach (left) and our embedding-based approach (right) for building generative models for connectomes. Both approaches incorporate neuronal features and statistical models based on them to induce a probability distribution over the space of networks. The standard approach requires feature engineering and handcrafted statistical models, while the embedding-based approach is agnostic to assumptions and available measurements of neuronal properties. **B**. A model architecture sketch of our implementation of the embedding-based approach: Latent neuronal features are modeled with an embedding layer, whose weights are tuned. The embeddings are fed to an MLP with a single hidden layer and a single output unit, which outputs the connection probability for the input neurons.

This approach may result in better models than those relying on explicit biophysical features, precisely by eliminating constraints that prior assumptions and biases impose, and may uncover features and relations that we did not know about or did not consider measuring. However, neither the superiority nor the interpretability of such models are guaranteed: Prior knowledge and intuition may prove critical inductive biases for building successful models; the optimization may converge to locally optimal or non-generalizing solutions; and even a successful model may remain opaque, making it computationally useful but biologically uninformative. Surprisingly, we show here that embedding-based models outperform biophysical feature-based ones in predicting neural connectivity, and, importantly, that the learned representations are interpretable — reflecting spatial and morphological properties of neurons and cell types as key organizational principles. This interpretability further enables a hybrid strategy, where recovered biological features anchor part of the representation while the remaining dimensions are left abstract, balancing predictive power with biological insight and offering a legible blueprint for cortical wiring.

## Results

### Learning neuronal features and wiring rules from connectivity data

To characterize the detailed circuitry within a cortical column of the primary visual cortex (V1) of a mouse [33, 34], we simultaneously learn an embedding for each neuron and train an ANN that estimates the connection probability of any two neurons, which we term the “Connector” (see Methods). Critically, to make these embeddings and the wiring rules interpretable, the Connector we train here is a small Multi-Layer Perceptron (MLP) [39, 40] with a single hidden layer. The input to the Connector is the learned embeddings of two neurons, and the learned input weights of each hidden unit in the Connector define a wiring rule on these abstract representations. Finally, the connection probability from one neuron to the other is given by a sigmoid function operating on the linear combination of all the wiring rules (Fig. 1B). Formally, the probability of a connection from neuron *i* to neuron *j* according to the model is

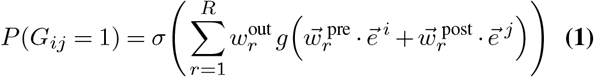

where 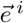 and 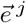 are the *D*-dimensional learned embeddings of the pre-synaptic and the post-synaptic neurons respectively, 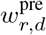 is the weight of the *d*-th coordinate of the embedding of the pre-synaptic neurons in the *r*-th wiring rule out of *R* rules (and similarly for 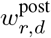 and the post-synaptic neuron), *g* is the Rectified Linear Unit (ReLU) function, 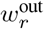 is the out-weight of the *r*-th wiring rule and 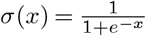 is the sigmoid function.

Individual connection probabilities across pairs of neurons can be organized into a matrix whose entry in row *i* and column *j* is *P*(*G*_*ij*_ = 1) (with *P*(*G*_*ii*_ = 0) ∀_*i*_). This is the average matrix induced by the model and we can use it to generate synthetic connectivity maps by sampling each connection according to the probability stored in the corresponding entry.

### Cortical connectome models that rely on a handful of embedding dimensions and wiring rules are highly accurate

We used this family of models to study the connectivity of 1351 neurons from a single cortical column in the primary visual cortex (V1) of a mouse [33, 34] (Fig. 2A). We binarize the connectome into a matrix *G* such that there is a synaptic connection from neuron *i* to neuron *j* (*G*_*ij*_ = 1) if at least one synapse was identified between them and are disconnected otherwise (*G*_*ij*_ = 0); we ignore auto-synapses (see Methods). We fit the model parameters, namely the Connector weights 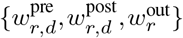 and the abstract learned neuronal features 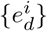, by maximizing the likelihood of our train data, and evaluate the model on test data, which are the result of a random split of neurons into two equally-sized sets (“train neurons” and “test neurons”). Importantly, our data split scheme is designed to address two key challenges: First, we must learn abstract embeddings for all neurons in the data to allow the inference of their connectivity patterns, and we need to hold out a subpopulation of neurons for evaluation, which is naturally defined by the subgraph over these test neurons. Second, to validate the learned wiring rules (the weights of the Connector 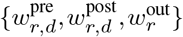), we need to show that they generalize to neurons that were not used for training. We therefore split neuronal pairs into 5 sets to train, validate, and test: (1) randomly chosen 80% of the pairs of train neurons (blue block in Fig. 2A) are used to learn the embeddings of the train neurons and the weights of the Connector (“train pairs”); (2) the remaining 20% (“validation pairs”) are used for convergence assessment and hyper-parameter tuning; (3) we then “freeze” the Connector, and learn only the embeddings of test neurons, using 80% of the connections between train neurons and test neurons (orange blocks in Fig. 2A; “used-mixed pairs”); (4) the remaining 20% of pairs of train neurons and test neurons (“validation-mixed pairs”) are not used for fitting parameters, but only to assess the convergence of the test neurons’ embeddings; (5) finally, the pairs of test neurons (black block in Fig. 2A) are the held out data that is used for evaluation of the model as a whole (“test pairs”).

**Fig. 2.**
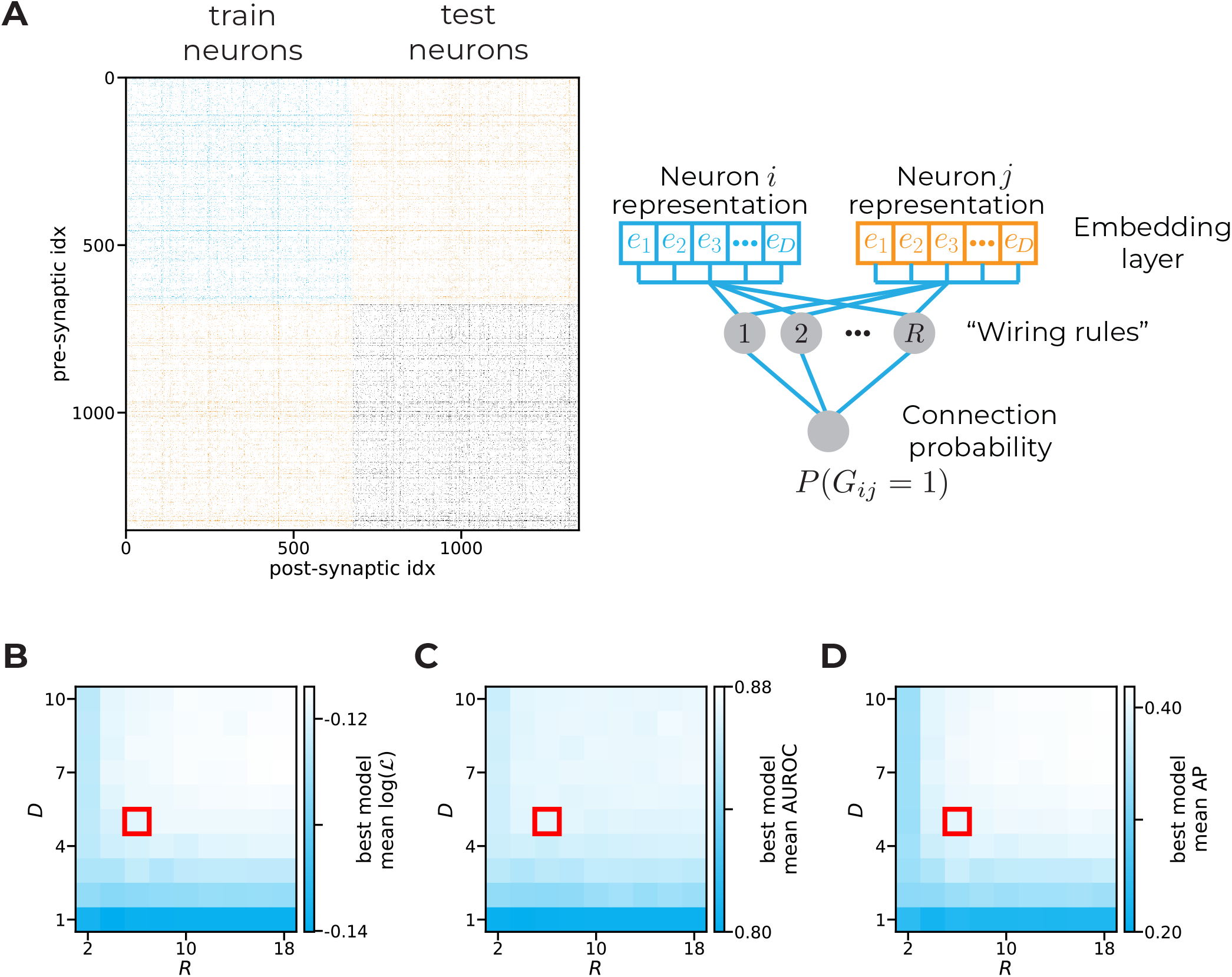
Five embedding dimensions and six wiring rules already saturate model performance on validation data. **A**. Left — the connectivity map of 1351 neurons from a cortical column in V1 of a mouse [33, 34]. Rows and columns in the matrix are sorted by the random split of neurons into train and test sets, and within a block are shuffled. Blocks in the matrix are colored by their part in the training and evaluation scheme. Right — a cartoon of the model’s architecture, as in Fig. 1B, where model parameters are colored by the blocks in the matrix to the left used to fit them (assuming neuron *i* is a train neuron and neuron *j* is a test neuron). The blue block (top left in the matrix, pairs of train neurons) is used to fit the embeddings of train neurons and the weights of the Connector. The orange blocks (top right and bottom left, pairs of train and test neurons) are used to fit the embeddings of the test neurons, given fixed fitted parameters for the rest of the model. The black block (bottom right, pairs of test neurons) is held out and used for evaluation. **B**. The mean log-likelihood over 3 data splits of the best hyperparameter configuration per embedding dimension (*D*) and hidden layer size (*R*) values. The red rectangle shows the chosen model dimensions, where performance saturates. **C**. Same as B for AUROC. **D**. Same as B for AP.

We trained models using multiple configurations of the models’ hyperparameters (see Methods) and compared them using several canonical classification metrics on the validation pairs: The log-likelihood, the Area Under the Receiver-Operating Characteristic Curve in predicting individual synaptic connections (AUROC), and the average precision (AP). Specifically, we explored how the performance of the models depends on the embedding dimension, *D*, and the number of units in the hidden layer or wiring rules of the Connector, *R*. Surprisingly, we find that performance saturates rapidly and that only 5 embedding dimensions and 6 wiring rules are sufficient to reach the full potential of this model architecture, regardless of the chosen metric (Fig. 2B-D, chosen dimensions are marked by red rectangles).

We therefore evaluate the performance of the model with 5 embedding dimensions and 6 wiring rules, using the held-out connectivity data between 675 test neurons, shown in Fig. 3A. The rows and columns are hierarchically sorted: First by excitatory (E) and inhibitory (I); within each of these two populations, they are sorted by their cortical layer; and within a layer by their classification into the 22 morphological types from [33, 34]. We emphasize that none of these properties were used in learning the embeddings or the wiring rules that predict the connectivity. We compare the embedding model, using a random data split for training and testing, with two variants of the “standard” generative models that we trained on a set of predefined biophysical features of neurons and handcrafted statistical formulations (as in Fig. 1A left). Specifically, we used the classification into morphological types ([33, 34]) to train a Stochastic Block Model (SBM) [41–43], and an Exponential Random Graph Model (ERGM) that uses these types and the physical distances between the somas of the neurons [31, 44, 45]. Notably, both SBM and the ERGM are the Maximum Entropy models over the space of networks, given their predefined set of features [31, 46], which means that these features are the “sufficient statistics” of these models. Thus, these are the unique generative models that conform to the observed values of the predefined features without making any additional assumptions: The SBM is the least-structured model that relies on the number of connections between all pairs of morphological types (which implies 22 × 22 features), and the ERGM adds to the constraints of SBM the sum of distances between connected neurons. Comparing sampled connectivity maps from the models we find that the SBM (Fig. 3B) is inferior to the ERGM (Fig. 3C), in terms of resemblance to the experimentally measured connectivity map (Fig. 3A). Importantly, the embedding model is even more accurate than the ERGM (Fig. 3D).

**Fig. 3.**
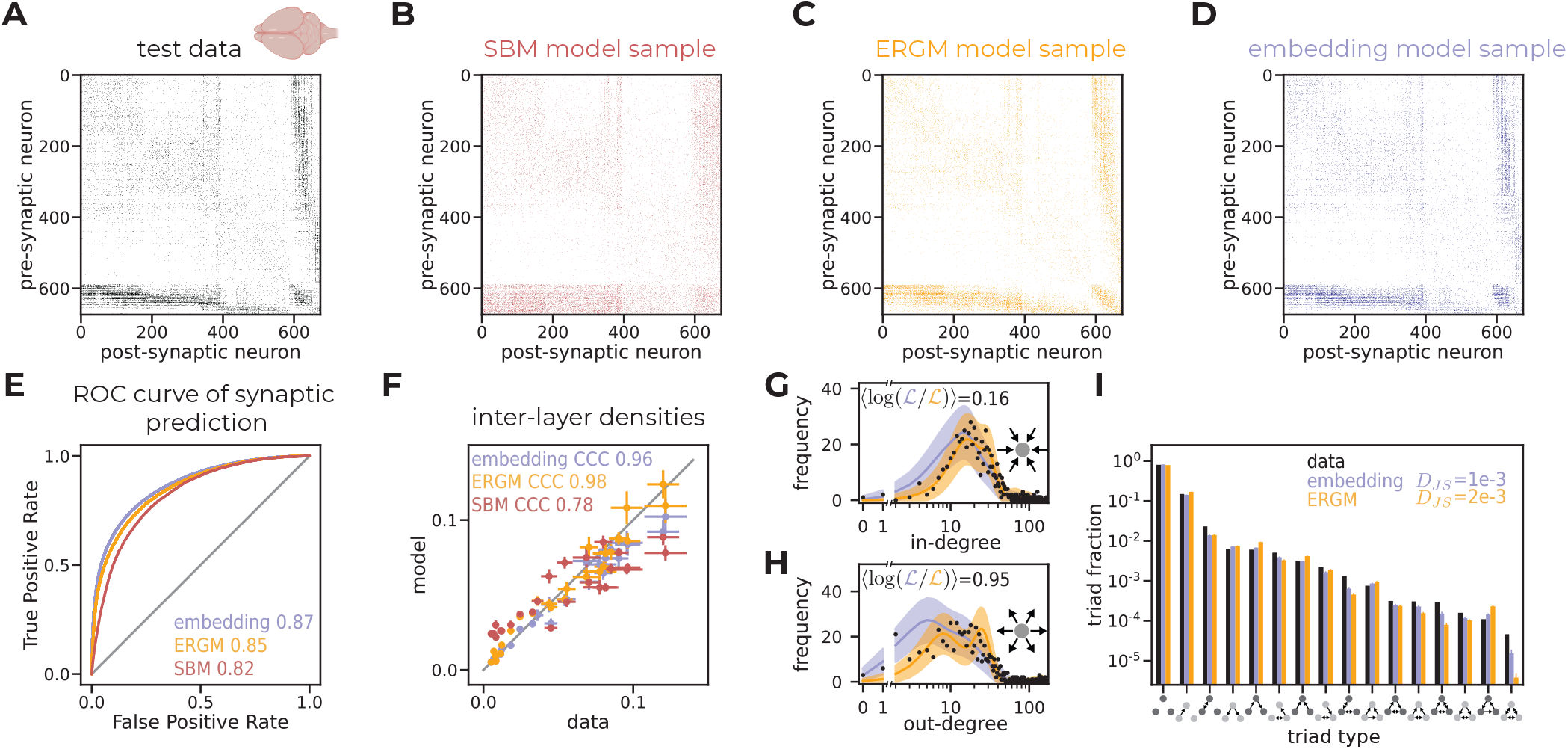
Models that rely on 5 embedding dimensions and 6 wiring rules are highly accurate. **A**. The connectivity map of the test data (held-out pairs of test neurons). Rows and columns are hierarchically sorted by E / I, within a type by layer, and within a layer by subtype (from [33, 34]). **B**. A synthetic connectivity map generated by the SBM. Rows and columns are sorted as in A. **C**. A synthetic connectivity map generated by the ERGM. Rows and columns are sorted as in A. **D**. A synthetic connectivity map generated by the embedding model. Rows and columns are sorted as in A. **E**. The ROC curves of synaptic prediction for the SBM (red), the ERGM (orange) and the embedding model (purple), calculated using the entries of the matrix from A as labels and those of the models’ average matrices as predictions. The area under each curve (AUROC) is specified in the legend. **F**. The densities of connections between cortical layers predicted by the SBM (red), the ERGM (orange) and the embedding model (purple) vs. the observed ones, for all pairs of layers. The *x*-coordinate shows the mean density over data bootstrapping (samples of subsets of neurons), and error bars show SD. The *y*-coordinate shows the mean of predicted average density by the model, for matching subset neuronal samples, and error bars show SD (see Methods). **G-H**. Degree distributions comparing the data (black), the ERGM (orange), and the embedding model (purple). Solid lines show average degree distributions, with shaded areas representing 2 SDs. The *x*-axis is linear from 0 to 1, then breaks and continues on log-scale. **G** shows in-degree (number of incoming connections) distribution. **H** shows out-degree (number of outgoing connections) distribution. **I**. The triplet motif distribution of the data (black), and the mean distribution of the ERGM (orange) and the embedding model (purple). Each bar shows the fraction of neuronal triplets out of all possible triplets that form a specific motif. There are 16 motifs up to isomorphism, hence 16 bars, which are sorted by their observed frequency in the data. Model bars show the mean over 100 samples, errorbars show 2 SDs.

We quantify the superiority of the embedding model by the Receiver-Operating Characteristic (ROC) curve, which summarizes the relation between the True Positive Rate (TPR) and False Positive Rate (FPR) of a binary classifier, and by the area under it (AUROC), which measures how well a binary classification model distinguishes between positive and negative examples. In our case, the ROC curve is based on the entries of the observed connectivity matrix from Fig. 3A as labels and those of the average matrices of different models as predictions. We find that the SBM achieves 0.824 ± 0.007, the ERGM reaches 0.849 ± 0.006, while our embedding model achieves 0.863 ± 0.007 (Fig. 3E, reported values refer to mean ± SD over 100 cross-validation splits). One-tailed t-tests revealed that the ranking is highly statistically significant (*p <* 10^−64^ for both pairs of models).

We further assess the models in predicting inter-layer cortical connectivity, which has been associated with information processing in the cortex [47–49]. We measure the agreement between the predicted inter-layer densities for all pairs of layers and the observed ones using the Concordance Correlation Coefficient (CCC) (see Methods). The SBM reaches a CCC value of 0.78 (bootstrap test, *H*_0_: CCC *>* 0.9, *p <* 10^−3^), while the ERGM and the embedding model reach 0.98 and 0.96, respectively (both with bootstrap test, *H*_0_: CCC ≤ 0.9, *p <* 10^−3^), reflecting highly accurate inter-layer connectivity predictions (Fig. 3F). Notably, however, the morphological types of excitatory neurons correspond to a classification of neurons into layers, thus the features used by type-based models carry information on layers by construction, whereas the embedding model does not rely on any information about layers. Both the ERGM and the embedding models recapitulate the connection tendency of individual neurons, reflected by the similarity of distributions of neuronal in-degree and out-degree values of the models to the observed in-degree and out-degree distributions (Fig. 3G-H). However, the embedding model is superior to the ERGM for neuronal degrees, as it assigns higher probabilities to the observed degrees (calculated like in [32]), which we quantify by the average log-likelihood ratio of individual degrees (over all neurons) that is positive for both in-degrees (0.16) and out-degrees (0.95). We further compared the observed distribution of small subcircuits or “network motifs” [50] with those predicted by the models. We find that the relative appearances of each of the 16 possible triplet motifs for the data and the embedding model are very similar (Fig. 3I), which we quantify by the Jensen-Shannon Divergence between the distributions, *D*_*JS*_ = 10^™3^ . This implies that one would need to observe a sample of ∼ 6.6×10^3^ triplets to be able to distinguish between the distributions with error rate of 1% [51]. Again, the ERGM shows inferior performance, compared to the embedding model, as it is less similar to the empirical distribution of motifs, with a *D*_*JS*_ value increasing to 2 × 10^™3^ (which implies that a two times smaller sample would suffice to meet the same error rate).

The superior performance and simplicity of the embedding model, compared to the types based ones, make it clear that this approach is preferable for learning generative models to describe the cortical connectivity. Furthermore, the fact that the embedding space and Connector that were tuned for the training set of neurons did so well in generalizing to successfully accommodate the held-out test neurons suggests that the coordinates in the latent space encode a “semantic” representation of neurons, whose “syntax” is defined and read by the Connector. We therefore turn to test the extent to which the embedding model has indeed inferred latent biological understanding of the space of neurons and wiring rules, beyond what it had been trained on — by presenting it with novel unseen biological settings.

### Embedding models generalize to predict connectivity of unseen cell types

To challenge our embedding models, we examine their ability to predict connectivity of neuronal types that were absent from the training set altogether: We train models using a split of the neurons into 2 halves for train and test, where all the neurons from three morphological types are only found in the test set. This setting simulates the scenario where the model is evaluated on new types, whose identities and connectivity between them were not available during training. Importantly, for type-based models, like the SBM or ERGM used above, where each pair of cell types has a dedicated parameter that captures the tendency of neurons of those types to connect, available connectivity data among all cell types is imperative for fitting all model parameters. We therefore compare our embedding models to a type-less baseline model that is agnostic of cell types, and incorporates only geometrical properties of neurons, namely their physical positions (see Methods).

We evaluate the models’ performance on the connections between the neurons of the triplet of held-out types they did not train on. Fig. 4A shows the observed connectivity map among 198 neurons of three randomly chosen types (one inhibitory and two excitatory), where rows and columns are hierarchically sorted by cell types and by layers of neurons. We find that the type-less baseline model clearly fails to predict the connectivity structure (Fig. 4B), and in particular misses especially dense connections of one of the classes (see the upper and left parts of the matrix). However, the embedding model captures the structure well (Fig. 4C). To reflect the nature of these differences in terms of the detailed circuit architecture, we compared the empirical distribution of triplet network motifs of the connectivity matrix between these three neuronal types and the motif distributions predicted by the models. The embedding model is very close to the empirical data and is clearly superior to the type-less baseline one, which misses the frequency of some motifs by orders of magnitude (Fig. 4D). We quantify the dissimilarity between the observed and predicted motif distributions with the Jensen-Shannon divergence, *D*_*JS*_, which shows a higher performance of the embedding model (0.02 compared to 0.07 for the type-less baseline model).

**Fig. 4.**
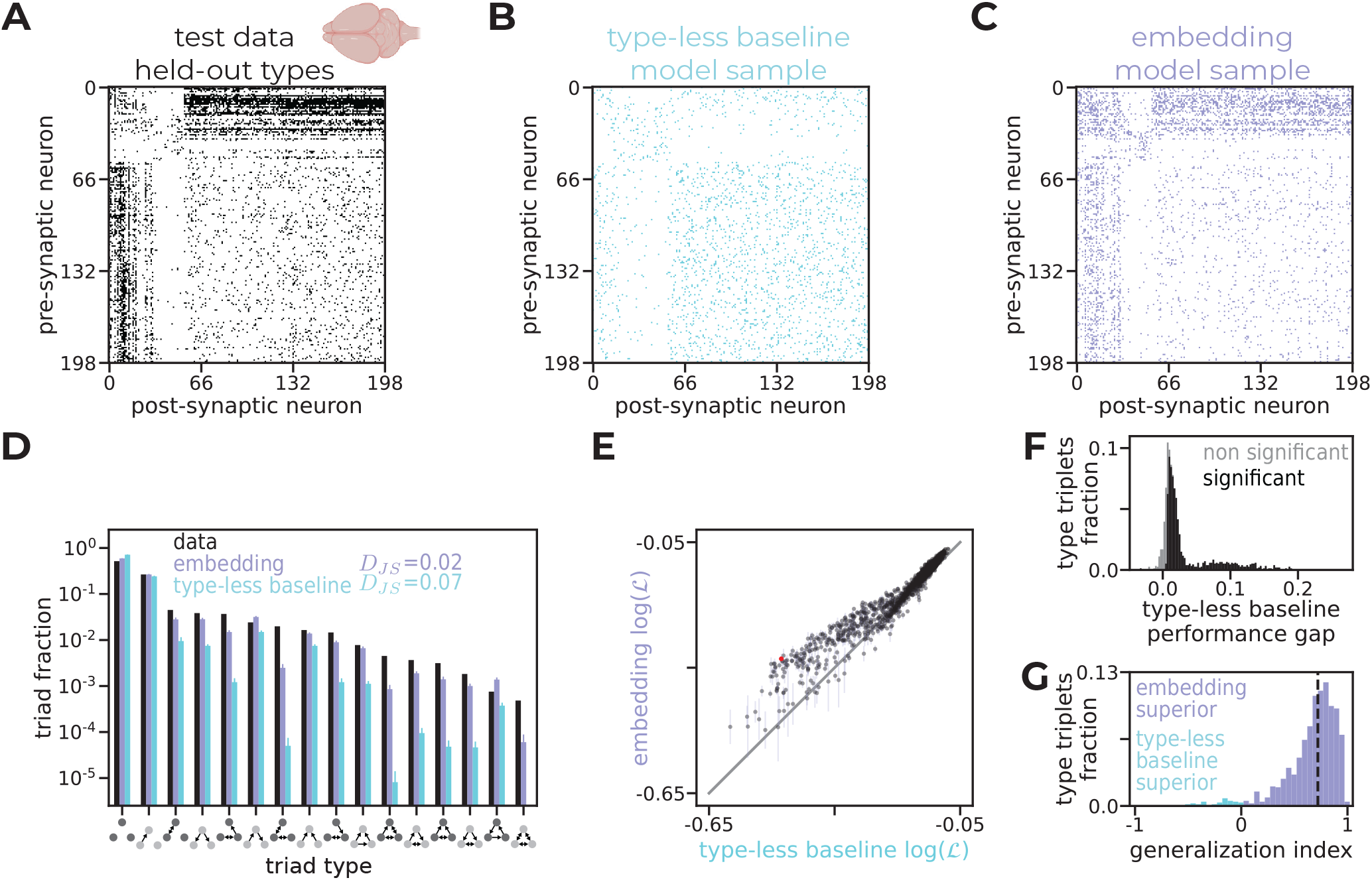
Embedding models can generalize to unseen cell types. **A**. The connectivity matrix between three randomly chosen cell types (one inhibitory - DTC and two excitatory - L2c and L3a) that were held out during training. Rows and columns are sorted by types. Inhibitory neurons are internally sorted by layer. **B**. A synthetic connectivity map between neurons of held out types, generated by the type-less baseline model. Rows and columns are sorted as in A. **C**. A synthetic connectivity map between neurons of held out types, generated by the embedding model. Rows and columns are sorted as in A. **D**. The triplet motif distribution of the data (black), and the mean distribution of the type-less baseline model (cyan) and the embedding model (purple) for the connectivity between the held-out cell types shown in panels A-C. Each bar shows the fraction of neuronal triplets out of all possible triplets that form a specific motif. There are 16 motifs up to isomorphism, hence 16 bars, which are sorted by their observed frequency in the data. Model bars show the mean over 100 samples, error bars show 2 SDs. **E**. Log-likelihood of the connectivity between held-out cell type triplets given by the embedding model vs. the type-less baseline model, over all 1540 triplets. Each dot represents one triplet, with coordinates showing the mean over 15 data splits in which all neurons of that triplet were held out. Error bars show SD (smaller than marker size on the *x*-axis). The red dot marks the triplet shown in panels A–C. The gray line is the identity. **F**. Distribution of performance gaps for the type-less baseline model (see main text) over all 1540 cell-type triplets. Bars are colored by the proportion of triplets with gaps that are statistically significant (dark, *p <* 0.01, permutation test, see Methods) or not (light) within each bar’s range. **G**. Distribution of the type triplet generalization index values, computed over the 1215 triplets with a statistically significant gap for which the full model is superior (panel F). For each triplet, the generalization index measures what fraction of the gap between the full embedding model and the baseline model is recovered by the embedding model with held-out types (see main text). Purple bars show triplets for which the embedding model with held-out types outperforms the baseline (positive values); cyan bars show the minority of triplets where the type-less baseline is superior (negative values). The dashed vertical line shows the median. Two extreme values of −1.7, −1.5 were trimmed for visibility.

To systematically compare the performance of the embedding model and type-less baseline one in the settings of unknown cell types, we perform this comparison for all 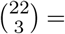 1540 different triplets of types and use 15 such data splits for each type triplet (where each split varies in the random assignment of the remaining neurons — those not belonging to the held-out triplet — to the training and test sets, while keeping the train-test ratio fixed). We quantified the performance of a model by the likelihood it gives to the observed connectivity map between the neurons of the held-out cell types. We find that the embedding model is better than the type-less baseline over many of the type triplets, and that the models show similar performance for some of the others (Fig. 4E).

To quantify the success of the models’ ability to generalize to neurons of new types, we define the “performance gap” of a model *M* for a type triplet *t* as the difference between the log-likelihood that the full embedding model that was trained using all cell types (from Fig. 3) gives to the connectivity map of a held-out type triplet *t* and the log-likelihood given by *M*: Δ_*M*_ (*t*) = *ℓ*_emb_full_(*t*) − *ℓ*_*M*_ (*t*). Fig. 4F shows the distribution of performance gap values over all held-out triplets for the type-less baseline model, Δ_base_: We find that this distribution has a large mode near zero and a heavy positive tail, which means that for many triplets of types, the full model and the type-less baseline that did not use these types in training give similar results. Thus, having access to these types does not add much to the model, suggesting that the sub-classification into types does not necessarily carry connectivity information.

We use a permutation test to filter triplets of types and focus only on those with a statistically significant performance gap (see Methods). For more than 99% of these 1221 triplets, the full model is superior to the type-less baseline, and for each of those, we calculate the “generalization index” of the embedding model, which we define as 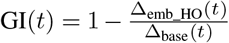. This quantifies the fraction of the gap between the full model and the type-less baseline model that is “closed” by the embedding model, which did not train on the held-out types. A value of 1 would indicate that it fully recovers the full model’s performance and 0 would indicate that it performs no better than the type-less baseline; negative values would indicate that the type-less baseline model is actually closer to the full model; values above 1 would indicate that the held-out embedding model outperforms the full model. We find that the embedding model that did not train on the held-out types successfully generalizes to predict their connectivity, as reflected by the histogram of generalization indices (Fig. 4G), which is heavily skewed towards 1 (median 0.72, interquartile range [0.57, 0.83]). Thus, in the realm of unknown cell types the embedding model is not only typically superior to a type-less baseline model that incorporates the physical layout of neurons, but is almost as good as the full model that used connectivity data among the hidden types for training.

We conclude that our embedding model generalizes to novel cell types, recovering most of the full model’s predictive accuracy without training on the connectivity of the held-out neurons. This implies that the latent representations learned by the model implicitly capture biological properties of the neurons that are substantial for determining the network’s structure. We proceed to ask whether we can uncover what these learned representations encode.

### Learned embeddings reveal biological features that predict connectivity

To understand what underlies the superior performance of the embedding models, we turned to explore the structure of the space of the learned neuronal embeddings, asking what the coordinates of neurons in that abstract space and the relations between them may reveal (see Fig. 5A). While these embeddings may be complex and highly non-linear, we find, surprisingly, that simply projecting the 5-dimensional representation of neurons into a 2-dimensional space using Principal Component Analysis (PCA) already reflects clear biological features: Coloring neurons in the 2-dimensional space according to their normalized cortical depth values revealed a global structure of the latent space that aligns with cortical depth (Fig. 5B, see Methods for the definition of normalized cortical depth). We quantify the correspondence between the coordinates of neurons in the latent space and cortical depth by the performance of a simple decoder in predicting cortical depth from the learned embeddings, which is highly accurate (*R*^2^ = 0.84 ± 0.04, see Methods). The global structure of the embedding space is further shaped by the Excitatory or Inhibitory identity of the neurons, which we found by projecting the 5-dimensional embeddings of neurons into a 2-dimensional space using PCA, but this time projecting neurons of different layers separately, which gave five such maps for layers 2-6. Coloring neurons in the low-dimensional space by their E / I identity reveals a clear separation of excitatory and inhibitory neurons in the latent embedding space (Fig. 5C shows neurons of layer 4 and Fig. SS1 the rest of the layers). We quantify this organization of the embedding space using a simple decoder that predicts neuronal E / I identities from learned embeddings, which reaches very high performance (*F*_1_ = 0.97 ± 0.01, see Methods). Our learned embeddings are also predictive of neuronal basal dendritic tree lengths, similarly reflected by the performance of a decoder that was trained to predict them (Fig. 5D *R*^2^ = 0.66 ± 0.07, see Methods). Together, these results show that the abstract embedding space that we learned is globally organized by biological features, and that these features are predictive of connectivity. Importantly, these results are robust and stable over cross-validation splits (Fig. SS2A-C). We emphasize that these biological features consistently emerge as candidates for shaping connectivity without prior assumptions, thanks to our computational framework.

**Fig. 5.**
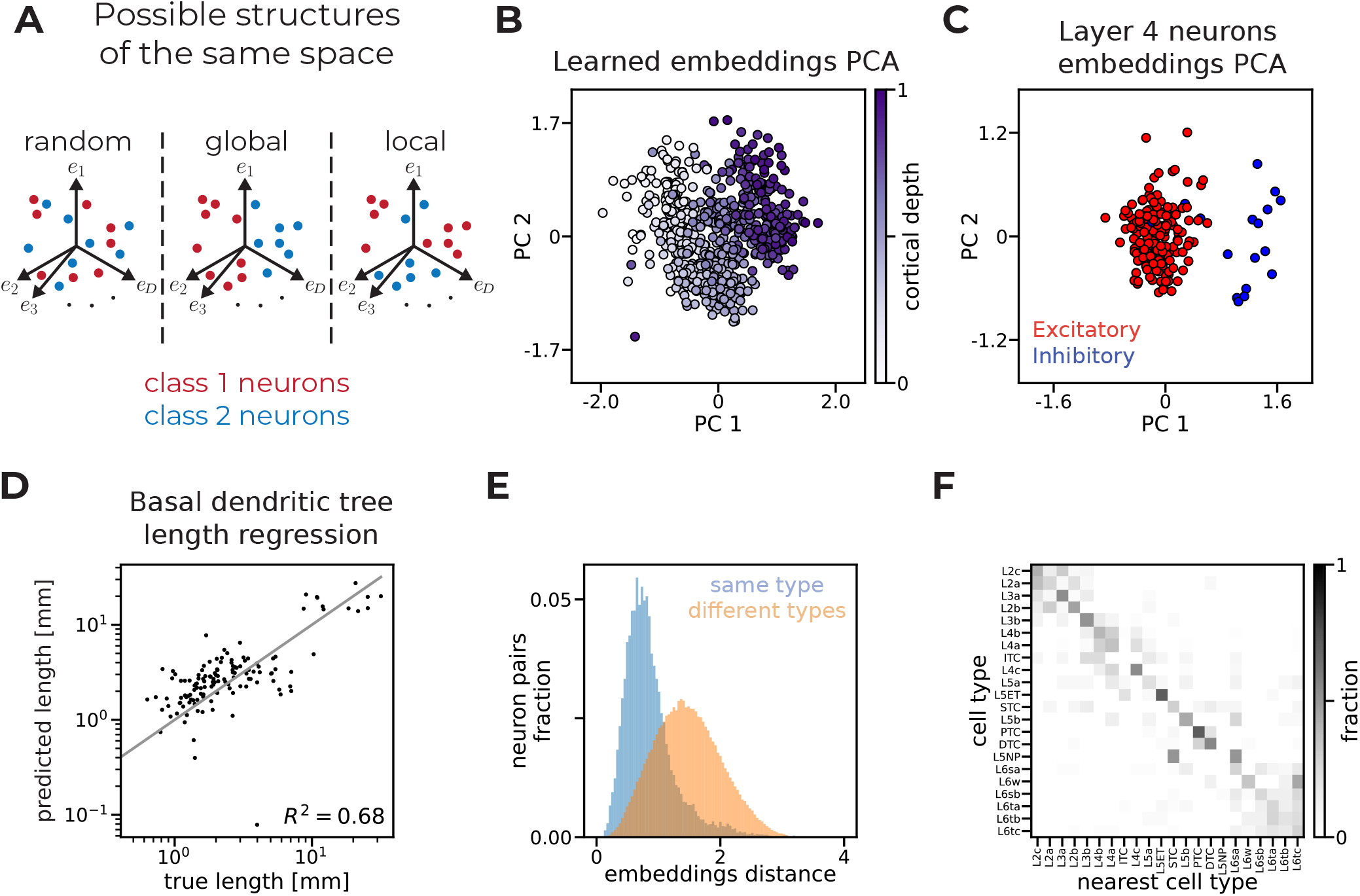
The learned abstract embedding space is shaped by biological features of neurons. **A**. A schematic illustration showing how neurons could organize within an abstract embedding space based on biophysical traits (such as neuron classes). The positions and distances between neurons in this latent space may convey information about various biophysical properties. **B**. The transformation of the learned embeddings into the plane of the first two Principle Components (PCs). Neurons are colored by their normalized cortical depth. **C**. The transformation of the learned embeddings of neurons of layer 4 into the plane of the first two PCs. Neurons are colored by their E / I identity. **D**. The prediction of basal dendritic tree length vs the true values. The predictor is an MLP with a single hidden layer with 10 units, which takes as inputs learned neuronal embeddings and outputs the basal dendritic tree length in millimeters. A predictor with median performance is shown (out of 10 trained predictors with different data splits, see Methods). Only data of held out neurons are scattered. **E**. The distributions of Euclidean distances in the learned abstract neuronal space of neurons of the same type (blue) and neurons of different types (orange), using the classification into 22 morphological types [34] **F**. The confusion matrix of a nearest-neighbor morphological type decoder. Rows are normalized such that they sum up to 1. Rows and columns are sorted to concentrate values near the main diagonal.

While we did not find a simple or immediate interpretation of the rest of the full 5-dimensional representation, we further examined the local structure of the learned embedding space, to uncover more “biological knowledge” it holds: We asked whether neighboring neurons in the latent space are similar in terms of their morphological types, which would reflect that the geometry of the embedding space holds fine semantic information about the neurons. We used, again, the classification of neurons in the data into 22 morphological types [34] and compared the Euclidean distances of neurons of the same type in our learned embedding space and neurons of different types. The distances between neurons of the same morphological type are much smaller than between neurons of different types (Fig. 5E, one-tailed t-test, *p <* 10^−300^). Furthermore, the nearest neighbor of a neuron in the embedding space is with high probability a neuron of the same morphological type, as shown by the “confusion matrix” in Fig. 5F, where the distribution of the types of the nearest cells are concentrated around the main diagonal (matrix values are normalized so that the rows sum up to 1). We quantify this by the accuracy of a nearest-neighbor morphological type decoder, which is 0.39, compared to a chance level of 0.09. This local organization indicates that biological features of neurons can be inferred from the features of their neighbors in the learned abstract space, that is based solely on connectivity data with no additional assumptions.

To verify that the morphological type information carried by the embeddings is not merely a reflection of the global structure by depth, E / I identity, and basal dendritic tree length uncovered above, we trained three classifiers for morphological types: One using only the learned embeddings, one using only the three biological features, and one using both. Evaluated over cross-validation splits, the embedding-only classifier (acc = 0.43 ± 0.03) and the biological-features-only classifier (acc = 0.54 ± 0.02) each outperform chance (acc = 0.09 ± 0.02). The gap is statistically significant (one-tailed t-test, *p <* 10^−100^ for both models). Additionally, the combined classifier achieves a higher accuracy than either alone (acc = 0.60 ± 0.02), which is again a statistically significant gap (one-tailed t-test, *p <* 10^−45^ for both models), demonstrating that the embeddings encode additional morphological type information that is not carried by the three biological features (see Fig. SS2D, Methods). This analysis proves that although we were unable to interpret the full 5-dimensional learned representations in clear biological terms, the embeddings encode additional biological properties of neurons, with predictive power for morphological cell types, that we cannot currently name. More comprehensive measurements of neuronal features would hopefully allow us to identify them.

### Inferring wiring rules’ logic and cortical design from the learned features of the embedding model

The accuracy of our models and the interpretability of some of its inferred features call for integrating them explicitly into the model and exploring the resulting wiring rules. We therefore learned a hybrid model, where neuronal representations are chimeric: Two of the coordinates stand for cortical depth and E / I identity that we inferred as key features and are fixed (not learned by the model), and the other three are learned embeddings as before (see Methods). We did not incorporate dendritic tree lengths into the hybrid representations since the cortical depth and E / I identities explain most of their variance, hence they add little information beyond the already used features. We use 6 wiring rules, which means that the model has the same dimensionality as earlier, but with fewer learned parameters. Importantly, this hybrid model is on par with the original embedding model (Fig. SS3). While we could not find an immediate biological interpretation of the other three dimensions of the learned representations of the hybrid model, identifying such neuronal traits may allow for reducing the dimensionality of the unknown features further. The compactness of the model means we can plot the complete and detailed architecture of the trained hybrid model, with 5-dimensional representation of each neuron and 6 wiring rules, where the edges in the Connector are colored according to their learned weights (Fig. 6A). The simplicity of this model means we can try to interpret the wiring rules in terms of the interplay between the features, asking which feature configurations drive each of the hidden units that implement the wiring rules to be active. We can then use the outgoing weights, that combine the hidden units’ activations to compute the connection probability in the output unit, to determine whether activation of each wiring rule is predictive of connectivity (positive weight) or lack thereof (negative weight).

**Fig. 6.**
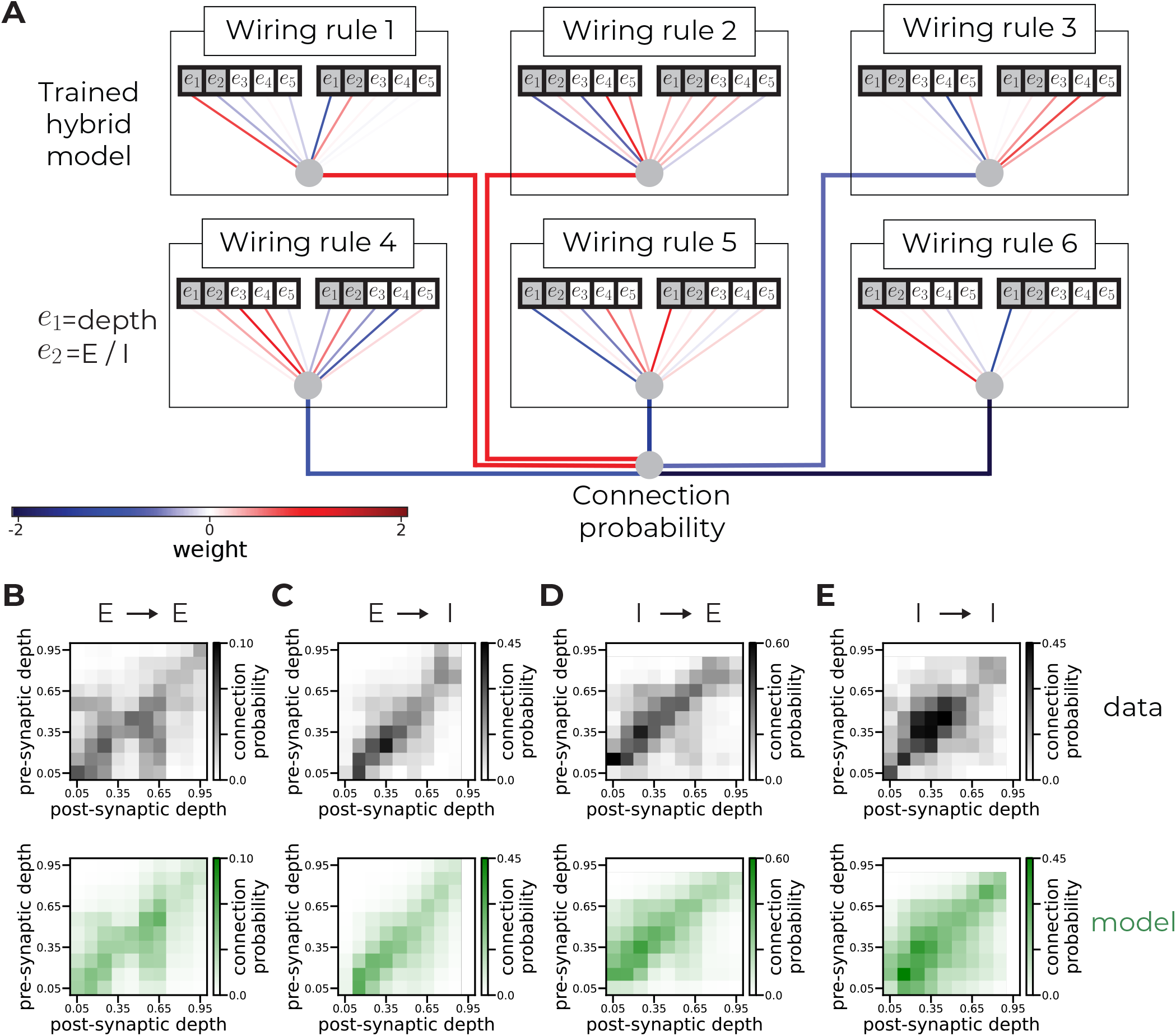
Hybrid models unlock wiring rules interpretability. **A**. The architecture of the trained hybrid model. Each neuron is represented by a 5-dimensional chimeric vector: The 2 first coordinates stand for cortical depth and E / I identity and are fixed (not learned); and the remaining 3 coordinated are abstract features learned by the model. Each hidden unit (wiring rule) receives the concatenation of a pair of neuronal representations as an input. The learned weights of the inputs are illustrated by edges between neuronal representations and the hidden units and are colored by their values. Neuronal representations were normalized to z-scores to compensate for different feature scales, thus the colors of the hidden weights correspond to z-scored inputs (see Methods). The out weights used to combine the activations of the hidden units to probability predictions are similarly illustrated as paths from the hidden units to the output node and colored by their learned weights. **B-E**. Connection probabilities as a function of pre-synaptic and post-synaptic neuron depths, conditioned on neuron types. Each panel shows empirical data (top) and corresponding model predictions (bottom). Depth ranges lacking neurons of a given type (and thus where connection probability is undefined) are colored white. This applies only to inhibitory neurons in the deepest bin (normalized cortical depths between 0.9 and 1.0). **B** excitatory to excitatory. **C** excitatory to inhibitory. **D** inhibitory to excitatory. **E** inhibitory to inhibitory.

For example, in rule 5 the weight of the depth of the presynaptic is negative and that of the post-synaptic is positive (and vice-versa for rule 6). Thus, rule 5 is “active” when the pre-synaptic is above the post-synaptic in the cortex, and rule 6 is active in the opposite scenario. As the absolute values of the weights of E / I identities are small, these rules are active for all E / I configurations (with a slight preference to excitatory neurons, as the weights are positive and we used 1 for E and − 1 for I). The outgoing weights of these rules are negative, which means that the predicted connection probability between neurons decreases when they are active, so we can interpret these rules as preventing neurons of different depths from connecting. While the other neuronal features take part in the computations carried out by rules 5 and 6, we can interpret the wiring logic in terms of the “marginal” over them by the above analysis. Accordingly, Fig. 6B-E shows the empirical and predicted connection probabilities based on the cortical depths, conditioned on the types of the pre-synaptic and the post-synaptic neurons: Indeed, the values in the matrices decrease when moving away from the diagonal, which is exactly increasing the difference of cortical depths. We validated this against the data by computing absolute cortical depth differences for connected and disconnected pairs: Disconnected pairs have larger differences, as expected, and a permutation test confirms this trend is significant (Fig. SS4A–B, *p <* 10^−4^, see Methods).

We further note that the weight of the cortical depth of the pre-synaptic neuron in rule 1 is positive, while that of the post-synaptic is negative. Thus, to activate rule 1, the pre-synaptic neuron has to be deeper than the post-synaptic one. Additionally, the weight of the E / I identity of the pre-synaptic neuron is negative and that of the post-synaptic neuron is positive, so rule 1 is active for inhibitory pre-synaptic neurons and excitatory post-synaptic ones. The outgoing weight is positive, so this rule predicts existence of connections. Together, inhibitory neurons tend to connect to excitatory neurons that are physically above them in the tissue, which aligns with the higher empirical probability values above the diagonal seen in the data (Fig. 6D top) and correctly predicted by the model as demonstrated by the similarity of the top and bottom panels in Fig. 6D. Permutation tests confirm that connected pairs of inhibitory and excitatory neurons have a significantly more positive mean depth difference than chance, and disconnected pairs a significantly more negative one (Fig. SS4C–D, both *p <* 10^−4^, see Methods).

Similarly, we can infer from rule 2 that excitatory neurons tend to connect to excitatory neurons below them, which also appears in empirical data: Probabilities below the diagonal are larger in the top panel of Fig. 6B. Again, this is correctly predicted by the model as demonstrated by the similarity of the top and bottom panels in Fig. 6B. Permutation tests confirm that connected pairs of excitatory neurons have a significantly more negative mean depth difference than chance, and disconnected pairs a significantly more positive one (Fig. SS4E–F, both *p <* 10^−4^, see Methods). Rule 3 deals mainly with the abstract features, so it cannot be interpreted in terms of depth and E / I identities, and rule 4 is an inversion of rule 2 (excitatory neurons tend to not connect to excitatory neurons above them, which is again observed by the asymmetric pattern of the matrices in Fig. 6B).

Together, the small set of wiring rules we have learned reveals simple organizational principles of cortical connectivity based on neuronal depth and E / I identities that can be summarized in words: Neurons with large depth differences tend not to connect, regardless of their type; inhibitory neurons tend to connect to excitatory neurons that are above them (physically) in the tissue; and excitatory to excitatory connections are preferred when the post-synaptic neuron is deeper than the pre-synaptic one. Although this is a partial sketch of how to wire up a cortical column using a subset of the features, it demonstrates how the simplicity of our model would allow one to delineate cortical wiring once more of the abstract features are unraveled.

## Discussion

Using a representational learning approach, we found that embedding neurons as points in a 5-dimensional space and using 6 simple wiring rules that act on these representations lead to accurate predictions of individual connections in a cortical column, empirical inter-layer connectivity, neuronal in-degrees and out-degrees, and small network motif distributions of a cortical column. We further showed that two dominant features of the abstract neuronal representations we have learned correspond to the cortical depth of a neuron and the Excitatory or Inhibitory identity of a neuron. Although we could not explain the other features of these embeddings in clear biological terms with the available measured neuronal properties, a hybrid model that uses these biological features and 3 additional abstract features is on par with the original model, and allowed us to give an outline for cortical wiring rules in biological terms. Importantly, the embeddings and the rules for cortical wiring turn out to be surprisingly simple and understandable. Interestingly, they do not require the detailed classification of tens of cell types, and moreover, the features and rules emerge from the model without any prior assumptions.

The three abstract features that we could not explain in immediate biological terms may imply that these abstract dimensions point towards biological and physical features that we do not currently measure, consider, or even have a clear name for. We expect that more data of both connectivity and neuronal characteristics could allow us to learn a mapping between the latent space and the measured biological space, and to identify the nature of the currently abstract dimensions. The ability of our abstract embedding space to predict the morphological types of neurons hints that our approach indeed offers a principled way to identify the key features without the need to guess or be biased by our current classifications or assumptions. In particular, future transcriptomic data at the level of individual neurons, aligned with their detailed connectivity, could help uncover the genetic and developmental interpretations of these features.

The approach we have taken here has focused on seeking neuronal embedding and a simple Connector, to allow for immediate and intuitive interpretation of the embeddings and rules, but a different choice of embeddings and nature of Connectors and their wiring rules could result in even more accurate and predictive models, or different interpretations of the features. On one extreme, seeking very simple wiring rules would require multiplicity of embedding dimensions and rich structure of the embedding space; an example of this would be lock-and-key mechanisms [52], which implement simple rules but require elaborate orchestration of chemical markers to produce complex structures. On the other extreme, very simple embeddings may require complicated non-linear connectivity rules that would weave these representations into a predictive model. Thus, characterizing the real features and rules that the cortex relies on requires more experimental data and measurements of various biophysical and genetic properties. However, the compactness and accuracy of the models we presented here already set a bound on the overall computational complexity necessary for building realistic connectomic structures of a cortical column, which is surprisingly low [53, 54].

Looking forward, our results present a general computational framework for studying the features and wiring rules that shape neural connectivity, and is immediately applicable to other datasets, brain areas, and species, and can be naturally extended for comparative analyses that can reveal conserved and variable design principles of neural systems [55]. Combined with neural activity data and simulations, they can serve as an infrastructure for delineating structurefunction relations in neural circuits [56, 57], and characterizing healthy and pathological conditions of circuit mis-wiring and activity [58].

## Methods

### Learned embeddings as parameterized linear projections

To obtain the learned embedding 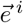 for neuron *i*, a 1-hot encoded vector 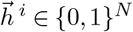 with a single 1 at position *i* and zeros elsewhere was passed through a linear embedding layer with learnable weight matrix **E** ∈ ℝ^*N* ×*D*^, where *N* is the total number of neurons and *D* is the embedding dimension:

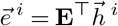

In this formulation, the embedding of neuron *i* corresponds directly to the *i*-th row of the transposed weight matrix, 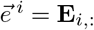. The embedding layer was initialized with Xavier uniform initialization and jointly optimized during training alongside the wiring rules of the Connector MLP.

### Connector architecture and training

The Connector was constructed as a small fully connected neural network with two layers. The input to the Connector consisted of the concatenated pre-synaptic and post-synaptic embeddings, 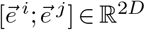. This concatenated vector was passed through a hidden layer of *R* units (the number of wiring rules), each computing a learned wiring rule. The output of the *r*-th hidden unit was given by a ReLU nonlinearity:

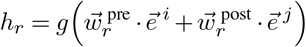

where 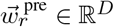 and 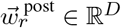 are the learned weights of the *r*-th wiring rule for pre-synaptic and post-synaptic embeddings, respectively. The hidden layer outputs were then combined via a final linear layer with learnable weights 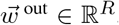, followed by a sigmoid function to produce a connection probability between 0 and 1. To ensure that the learned wiring rules operate directly on the embedding representations without bias degrees of freedom, the models do not include biases.

Weights in the hidden layer (matrix **W**^pre^ and **W**^post^ collectively, and output weight vector 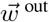) were initialized with Xavier uniform initialization and updated via backpropagation. The embedding layer parameters were jointly learned with the Connector weights during training.

### Data pre-processing

The connectome is based on [33, 34] and consists of 1351 neurons. Connections are binarized, namely a pair of neurons is connected if and only if at least one synapse exists between them, and auto-synapses (connections from a neuron to itself) are ignored, resulting in 75251 connections. Data (connectivity, neuronal positions, and cell types) are retrieved using CAVE (Connectome Annotation Versioning Engine) [59], and based on data release 1507 (from July 2025). Cell types are based on the allen_column_mtypes_v2 table. The layer of excitatory neurons is determined by their subtype (that is layer-based), and of inhibitory neurons, by the layer of the excitatory neuron with the vertically closest soma. To normalize cortical depths, the vertical position of each neuron, *v*, is transformed to a value between 0 and 1 using the linear transformation 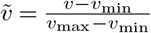, where *v*_min_ and *v*_max_ denote the minimal and maximal values in the data, respectively. The normalized depth is then 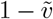. Neuronal basal dendritic tree lengths are retrieved from the public BossDB repository (S3 bucket `bossdb-open-data`, prefix `iarpa_microns/minnie/minnie65/skeletons/v661/metadata`).

### Model hyperparameters

The hyperparameters of the model were tuned with a grid search. Specifically, the following hyperparameters and options were considered: 10 initialization seeds; 10 embedding dimensions, ranging from 1 to 10; 9 hidden layer sizes, with values in {2*i* | 1 ≤ *i* ≤ 9, ∈ N}; 4 learning rates, with values in {0.001, 0.003, 0.1, 0.3}; 4 regularization coefficients for the learned neuronal embeddings, with values in 0.3, 0.1, 3, 10 ; same 4 regularization coefficients for the Connector weights; 3 data splits (dividing the neurons into 2 halves of train and test randomly, see main text). This resulted in 172800 hyperparameter configurations. All configurations were trained using the Adam optimizer, with a batch size of 64 for 200 epochs. Data were randomly split into 80% train and 20% validation samples and for each hyperparameter configuration, the model with the best validation loss over all epochs was chosen. Then, the validation loss on the train neurons was averaged for each hyperparameter configuration up to the data splitting index (3 values were averaged for each configuration), and the one with the minimal loss was chosen. As performance saturated rapidly with model dimensionality (see main text), the dimensions for the final hyperparameter configuration were fixed in advance, and only configurations with 5 embedding dimensions and 6 hidden units were considered. All visualizations use the first data split. To gain statistical power for cross-validation splits, 1000 additional models were trained using the chosen hyperparameter configuration — using 100 cross-validation splits and 10 initialization per split. Then, the best model per split (out of the 10 initializations) was chosen, resulting in 100 cross-validation splits using the chosen hyperparameter configuration.

### Inter-layer cortical connectivity

The density of connections between neurons of layers *m, k* in graph *G* is given by

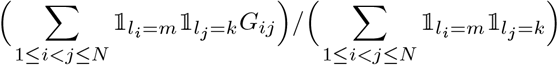

where *N* is the number of neurons in the network, *l*_*i*_ is the layer of the neuron *i*, 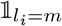 is an indicator that takes 1 if the neuron *i* is in the layer *m* and 0 otherwise, and *G* is the connectivity matrix. The empirical inter-layer density was obtained by substituting the observed adjacency matrix for *G*; the mean density predicted by the model was obtained by substituting the model’s average connectivity matrix for *G*. Agreement between predicted and observed inter-layer densities across all pairs of layers was quantified using the Concordance Correlation Coefficient (CCC), a measure of agreement between continuous variables bounded in [− 1, 1], where a value of 1 indicates perfect agreement. A bootstrap procedure was used to construct a sampling distribution of the CCC under each model. At each of 1000 bootstrap iterations, a random subset of 75% of neurons was drawn without replacement, and the subnetwork induced by that subset was used to compute the inter-layer density for both the data and the model. The CCC between the predicted and observed densities was then computed for each subsample, yielding a distribution of 1000 CCC values per model. One-sided bootstrap tests were performed to evaluate model accuracy against a threshold of CCC_0_ = 0.9, chosen as a standard criterion for strong agreement. For the SBM, the null hypothesis *H*_0_: CCC *>* 0.9 was tested, and the *p*-value was computed as the proportion of bootstrap samples in which the CCC exceeded 0.9. For the ERGM and embedding model, the null hypothesis *H*_0_: CCC ≤ 0.9 was tested, and the *p*-value was computed as the proportion of bootstrap samples in which the CCC fell at or below 0.9.

### A baseline model for generalizing to unseen cell types

Since type based models (like SBMs and ERGMs with type-based features) have a dedicated parameter to each pair of cell type that captures their tendency to connect, connectivity data between all cell types is required to learn all model parameters. Thus, the model considered as a baseline for learning with held-out types, is an ERGM that relies on 2 features — the total number of connections and the sum of distances between connected neurons. This is a leaner version of an ERGM that learns to recapitulate only the overall connectivity and its spatial properties, but is agnostic to type-specific structures.

### Identifying cell-type-specific performance gaps between models

To assess whether the full embedding model and the baseline model with held-out types differ in their ability to predict type-conditioned connectivity, a permutation test was applied separately for each cell type triplet.

For each of the 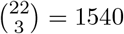 triplets of held-out cell types, the log-likelihood of each model on the connections between neurons of the held-out types in the test set was computed over independent cross-validation splits, yielding a scalar performance measure per split. The baseline model was evaluated using 15 cross-validation seeds, while the full embedding model was evaluated using 100 seeds, resulting in 15 log-likelihood values per triplet for the baseline and 100 values for the full embedding model.

To determine whether the two models exhibited a statistically significant performance gap for a given triplet, a two-tailed permutation test was performed. The observed test statistic was defined as the difference in mean log-likelihood between the full embedding model and the baseline model. Under the null hypothesis that both sets of values are drawn from the same distribution, the combined pool of 15 + 100 = 115 values was permuted, and at each permutation a new mean difference was computed by randomly reassigning values to groups of sizes 15 and 100. This procedure was repeated for 10,000 permutations, and the *p*-value was computed as the proportion of permutations in which the absolute permuted difference was greater or equal than the absolute observed difference. A significance threshold of *α* = 0.01 was used.

### Decode biophysical features from embeddings

The biophysical properties of neurons were decoded from the 5-dimensional embedding vectors of test neurons. For all decoders, performance was evaluated using 10-fold cross-validation with random shuffled splits, each holding out 20% of the neurons as a test set. The metrics reported in the main text refer to the mean ± SD on the held-out neurons over this cross-validation, unless specified otherwise.

### Cortical depth

A polynomial regression model of degree 2 was used to predict the normalized cortical depth (see above) of each neuron from its learned embedding. Performance is reported as the coefficient of determination (*R*^2^).

### Excitatory / Inhibitory identity

A logistic regression classifier was trained to predict the excitatory or inhibitory identity of each neuron from its learned embedding, where excitatory neurons were labeled as 1 and inhibitory neurons as 0. Performance is reported as the *F*_1_ score.

### Basal dendritic tree length

An MLP regressor with a single hidden layer of 10 units was used to predict the basal dendritic tree length of each neuron from its learned embedding. Performance is reported as the coefficient of determination (*R*^2^).

### Morphological type classification

To assess whether the learned embeddings encode morphological type information beyond what is captured by the three biophysical features above, three Random Forest classifiers (100 trees each) were compared. The first classifier was trained on the standardized biophysical features alone (cortical depth, E / I identity, and basal dendritic tree length). The second was trained on the 5-dimensional embedding vector alone. The third was trained on the concatenation of both feature sets. Performance is reported as the accuracy. The metrics reported in the main text for these classifiers are the mean ± SD over 100 cross-validation splits of neurons into train and test (see hyperparameters subsection). The metric value for each split was taken as the mean over 10-fold cross-validation of the test neurons of that split.

### Hybrid models

In the hybrid model, the chimeric neuronal representations are 5-dimensional vectors:

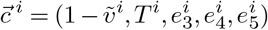

where the first 2 coordinates are fixed (not learned) and correspond to biophysical features of neurons: Normalized cortical depth (denoted by 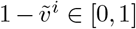, see above) and Excitatory (E) or Inhibitory (I) identity (encoded as 1 and − 1, respectively and denoted by *T* ^*i*^). The remaining 3 coordinates, 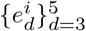, are the 3 learned abstract dimensions.

The model uses 6 hidden units, implementing wiring rules based on the chimeric representation. Together, following Eq. (1), the probability predicted by the model is given by

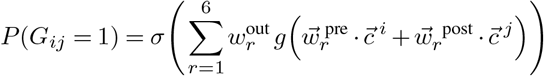

The hybrid model is trained with a single hyper-parameter configuration that is chosen to be the best performing one from the sweep done to tune the hyper-parameters of the original model. The same data splitting strategy and sequential training pipeline were used to train it.

### Transform model embeddings to z-score

The *d*-th coordinate of the z-scored embedding of neuron *i* is defined by

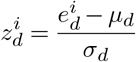

where 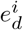 is the *d*-th coordinate of the embedding of neuron *i*

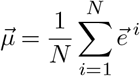

is the mean embedding (denoting the number of neurons by *N*) and

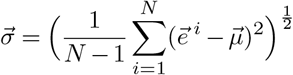

is the standard deviation of embeddings along each axis. To compensate for model embeddings transformed to z-scores, the hidden weights of the model are multiplied by 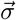 entry-wise, such that the new weights of the *r*-th wiring rule become (using the notation from Eq. (1))

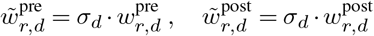

and a bias vector 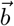 is added to the hidden units, whose *r*-th coordinate is defined by

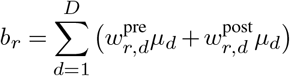

The output of the *r*-th hidden unit for input neurons *i, j* is thus given by (adapted from Eq. (1))

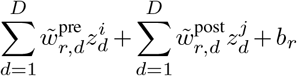

which equals the output of the same unit for the same neurons when performing inference with the original model.

### Analysis of cortical depth differences between connected and disconnected neuron pairs

To assess whether connected neuron pairs exhibit systematically different cortical depth relationships than unconnected pairs, the signed depth difference Δ*d*_*ij*_ = *d*_*i*_ − *d*_*j*_ was computed for each ordered pair (*i, j*), where 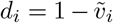 denotes the normalized cortical depth of neuron *i* and 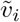 is its normalized vertical position. This quantity was computed separately for all pairs sharing a directed synaptic connection (connected pairs, indexed by *C*) and for all ordered pairs with no connection (disconnected pairs, indexed by *D*), with the analysis optionally restricted to pairs whose pre-synaptic and post-synaptic neurons belong to specified cell types. For the All → All analysis (namely, considering both excitatory and inhibitory neuron pairs), the absolute depth difference | Δ*d*_*ij*_ | was used in place of the signed quantity. The observed means of the empirical distributions of Δ*d*_*ij*_ for connected and disconnected pairs are given by

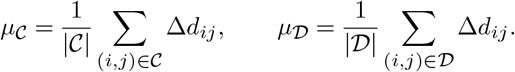

To determine whether *µ*_C_ and *µ*_D_ deviated significantly from chance, a permutation test was applied. The combined pool of *N* = |C| + |D| depth differences was randomly partitioned into a surrogate connected group of size |C| and a surrogate disconnected group of size |D|, yielding surrogate means 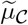 and 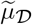. This procedure was repeated for 10,000 permutations, producing null distributions of 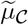 and 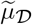. Directional *p*-values were then computed as

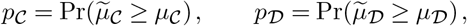

where the inequality is reversed for tests directed toward a more negative observed mean, and the probabilities are taken over permutations. For *p*-values of 0 we used 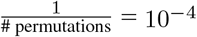 as an upper bound.

## ACKNOWLEDGMENTS

We thank members of Schneidman’s lab for critical suggestions and insights. This work was supported by Simons Collaboration on the Global Brain grant 542997, Israel Science Foundation grant 137628, Azrieli Institute for Brain and Neural Sciences and the Hedda, Alberto, and David Milman Baron Center for Research on the Development of Neural Networks of the Weizmann institute, as well as the Knell family Institute for Artificial Intelligence, Martin Kushner Schnur, and Mr. & Mrs. Lawrence Feis. ES is the incumbent of the Joseph and Bessie Feinberg Chair.

## Supplementary figures

**Fig. S1.**
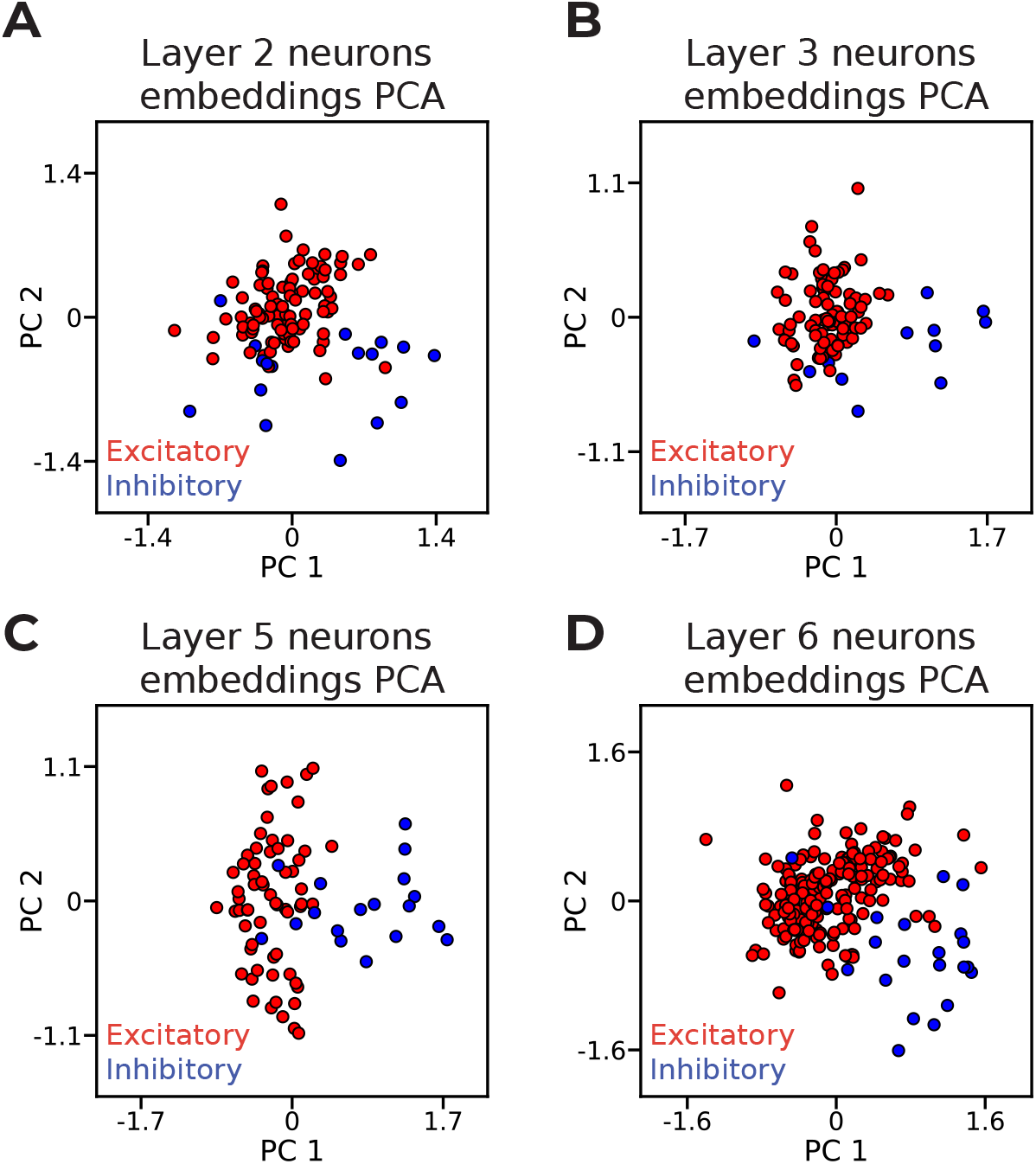
The learned abstract embedding space is shaped by neuronal E / I identity. A-D. The transformations of the learned embeddings of neurons of each layer into the plane of the first two PCs. Neurons are colored by their E / I identity. A — neurons of layer 2; B — neurons of layer 3; C — neurons of layer 5; D — neurons of layer 6.

**Fig. S2.**
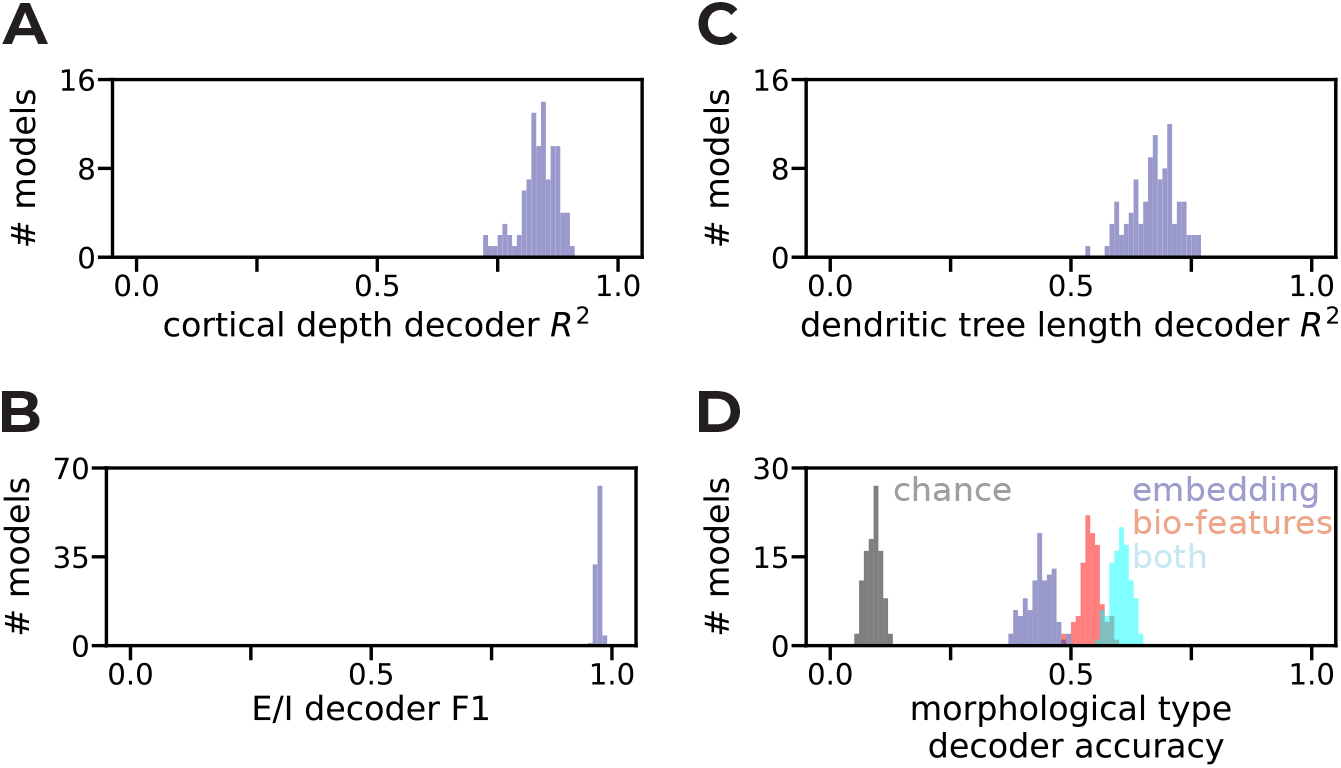
Neuronal biophysical features are consistently decodable from learned embeddings. **A-C**. The performance distribution of biophysical features decoders from learned embeddings over 100 data splits. A — cortical depth regressor. B — Excitatory/Inhibitory identity regressor. C — basal dendritic tree length regressor. **D**. Accuracy of three random forest classifiers for morphological types over 100 data splits: Using only the learned embeddings (purple), using only the three biological features decoded in A-C (pink), using both (cyan), and chance level (gray). See also Methods.

**Fig. S3.**
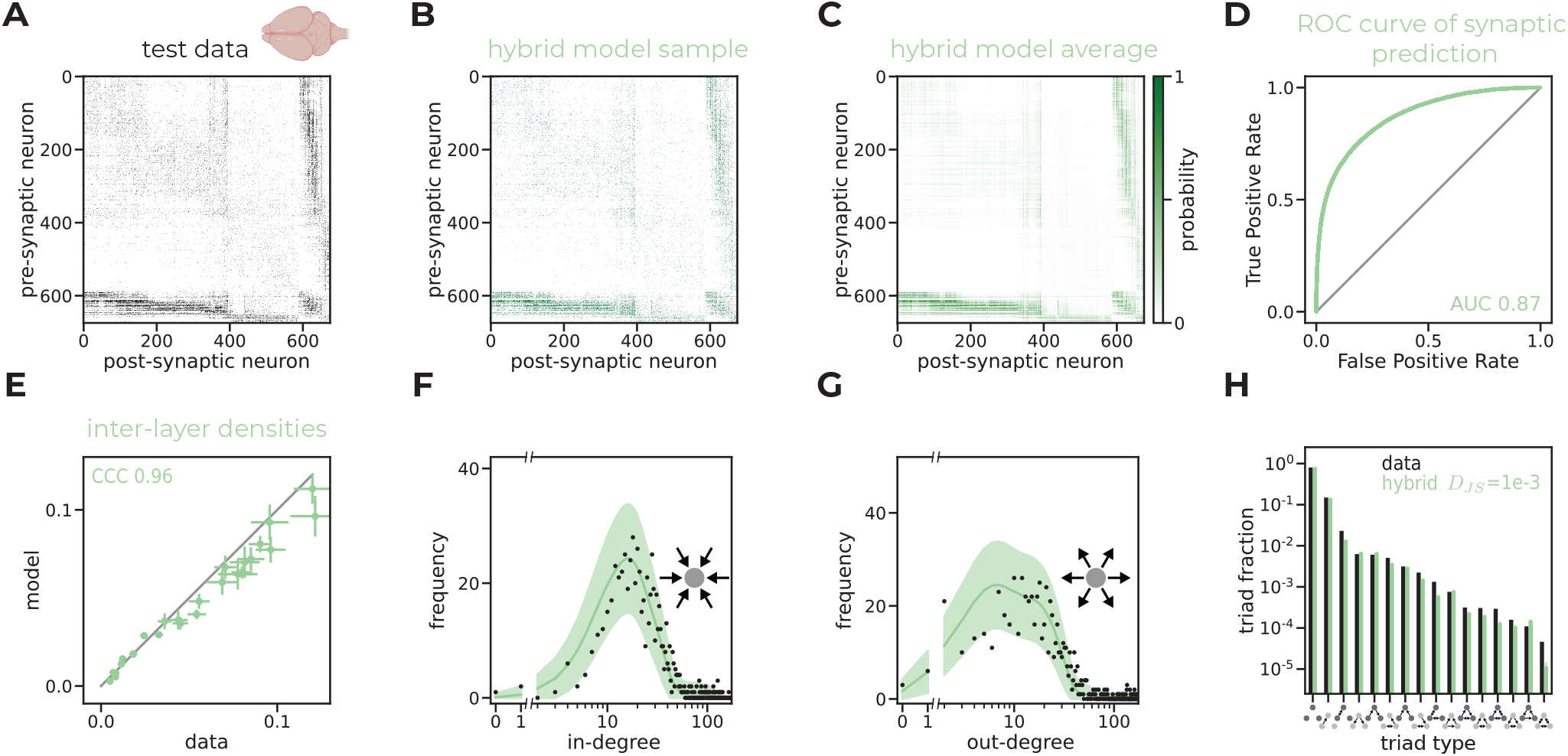
The hybrid model performs on par with the original model in predicting connectivity. **A**. The connectivity map of the test data (held-out pairs of test neurons). Rows and columns are sorted by E / I, within a type by layer, and within a layer by subtype (from [33, 34]). **B**. A synthetic connectivity map generated by the hybrid model. Rows and columns are sorted as in A. **C**. The average connectivity map of the hybrid model (namely, the connection probability it assigns to each pair of neurons). Rows and columns are sorted as in A. **D**. The ROC curve of synaptic prediction, calculated using the entries of the matrix from A as labels and those of the matrix from C as predictions. **E**. The densities of connections between cortical layers predicted by the hybrid model vs. the observed ones, for all pairs of layers. The *x*-coordinate shows the mean density over data bootstrapping (samples of subsets of neurons), and error bars show SD. The *y*-coordinate shows the mean of predicted average density by the model, for matching subset neuronal samples, and error bars show SD (see Methods). **F-G**. Degree distributions comparing the data (black) and the hybrid model (green). Solid lines show average degree distributions, with shaded areas representing 2 SDs. The *x*-axis is linear from 0 to 1, then breaks and continues on log-scale. **F** shows in-degree (number of incoming connections) distribution. **G** shows out-degree (number of outgoing connections) distribution. **H**. The triplet motif distribution of the data (black) and the mean distribution of the hybrid model (green). Each bar shows the fraction of neuronal triplets out of all possible triplets that form a specific motif. There are 16 motifs up to isomorphism, hence 16 bars, which are sorted by their observed frequency in the data. Model bars show the mean over 100 samples, errorbars show 2 SDs.

**Fig. S4.**
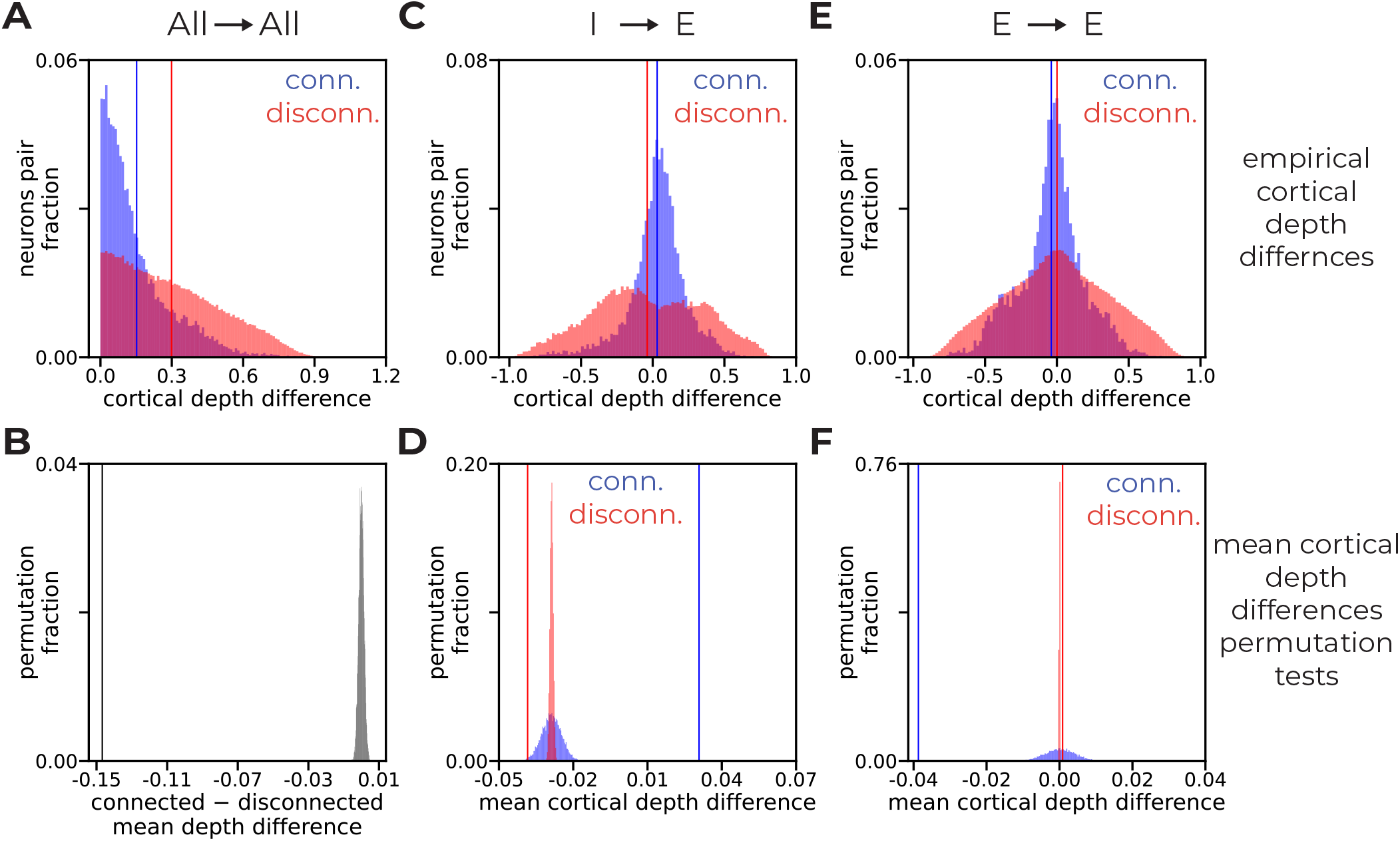
Connected neuron pairs exhibit structured cortical depth differences. **A, C, E**. Distributions of cortical depth differences for connected (blue) and disconnected (red) neuron pairs. **A**. Absolute values of cortical depth differences (|pre-synaptic depth − post-synaptic depth|) for all neuron type pairs. **C**. Signed cortical depth differences (pre-synaptic depth minus post-synaptic depth) for inhibitory-to-excitatory pairs. **E**. Signed cortical depth differences for excitatory-to-excitatory pairs. Vertical lines indicate the mean of each distribution. Positive signed values indicate the pre-synaptic neuron is deeper than the post-synaptic neuron. **B**. Null distribution of the difference between connected and disconnected mean absolute depth differences under a permutation test (All → All), with the observed value indicated by a vertical line. **D, F**. Null distributions of connected (blue) and disconnected (red) mean depth differences under permutation tests, for Inhibitory → Excitatory (**D**) and Excitatory → Excitatory (**F**) pairs. Vertical lines indicate the observed means. See also Methods.

## Notes

### Competing Interest Statement

The authors have declared no competing interest.

